# Structural basis of RNA polymerase recycling by the Swi2/Snf2 ATPase RapA in *Escherichia coli*

**DOI:** 10.1101/2021.05.05.442812

**Authors:** M. Zuhaib Qayyum, Vadim Molodtsov, Andrew Renda, Katsuhiko S. Murakami

## Abstract

After transcription termination, cellular RNA polymerases (RNAPs) are occasionally trapped on DNA, impounded in an undefined Post-Termination Complex (PTC), limiting free RNAP pool and making transcription inefficient. In *Escherichia coli*, a Swi2/Snf2 ATPase RapA is involved in countering such inefficiency through RNAP recycling. To understand its mechanism of RNAP recycling, we have determined the cryo-electron microscopy (cryo-EM) structures of two sets of *E. coli* RapA-RNAP complexes along with RNAP core enzyme and elongation complex (EC). The structures revealed the large conformational changes of RNAP and RapA upon their association implicated in the hindrance in PTC formation. Our study reveals that although RapA binds away from the DNA binding channel of RNAP, it can close the RNAP clamp allosterically thereby preventing its non-specific DNA binding. Together with DNA binding assays, we propose that RapA acts as a guardian of RNAP by which prevents non-specific DNA binding of RNAP without affecting the sigma factor binding to RNAP core enzyme, thereby enhancing RNAP recycling.

## INTRODUCTION

Bacterial multi-subunit DNA-dependent RNA polymerase (RNAP) is a fast and processive enzyme that can perform RNA synthesis at a rate of 90 nt/sec for thousands of nucleotides during elongation phase of transcription in a purified system (1). However, in biological systems, this process is not monotonous but encompasses a wide array of regulatory mechanisms giving different functional outputs. Transcriptional pausing is one such process and plays important roles in transcription regulation, RNA folding and transcription-translation coupling (2,3). It also serves as an early step during transcription termination (4-7). A transcribing RNAP also halts when it encounters a DNA lesion (8). Therefore, the transcribing RNAP requires a series of accessary factors to achieve transcription speeds like those achieved in a purified system.

General elongation factors NusA and NusG bind RNAP to regulate the rate of transcription, half-life of transcription pausing, mediate transcription-translation coupling, and facilitate transcription termination (9-14). Due to nucleotide misincorporation into RNA, a backward translocation of RNAP along DNA (as known as RNAP backtracking) temporarily arrests RNA synthesis (15,16). To mitigate such events, transcription factors GreA/B bind to RNAP secondary channel to cleave backtracked RNA including mis-incorporated nucleotides for restarting RNA synthesis (17,18).

DNA damage, stalling events, and collisions with template-bound proteins halt RNA elongation. Mfd, an ATP-dependent motor enzyme, is a part of transcription-coupled repair (TCR) system that binds upstream DNA to the stalled RNAP, triggers transcription bubble collapse, and the dissociation of RNAP from the DNA by virtue of its translocase activity (17,19-21). A recent study also reveals the participation of an RNAse (RNAse J1) in resolving such stalled RNAPs (22). The last phase of the transcription cycle (as known as transcription termination) is also regulated by another ATP-dependent helicase/translocase Rho, that translocates along nascent RNA and dislodges the EC after interacting with RNAP, NusA and NusG (23-26).

During the transcription termination, RNAP releases RNA product and dissociates from the template DNA to prepare for a next round of transcription (RNAP recycling). However, some RNAPs form an undefined post-transcription/post-termination complex (PTC) (27) that prevents RNAP recycling, thus inhibits gene expression. In fermicutes and actinobacteria, transcription factor HelD, a RNAP associated <u>s</u>uper<u>f</u>amily 1 (SF1) ATPase, is involved in RNAP recycling (28,29). Recent cryo-EM studies of the RNAP-HelD complex (29-31) revealed that the HelD inserts its two domains deep into the RNAP main and secondary channels akin a pronge to de-stabilize the EC. Dissociation of HelD from the RNAP is an active process that utilizes energy from ATP hydrolysis by HelD (29-31).

In proteobacteria including *E. coli*, RNAP recycling is facilitated by RapA, a Swi/Snf2 protein superfamily ATPase (32) (Fig. 1A). RapA was first observed as a co-purifying protein named τ factor with *E. coli* RNAP about 50 years ago in a study that discovered promoter specificity σ^70^ factor (33). The studies on Swi2/Snf2 family of enzymes have mostly focused on their roles in chromatin and nucleosome remodeling in eukaryotes (34). Interestingly, some members of the family, such as RapA, do not modify chromatin structure. Instead, its direct interaction with RNAP enhances RNA expression by facilitating RNAP recycling (35).

**Figure 1.**
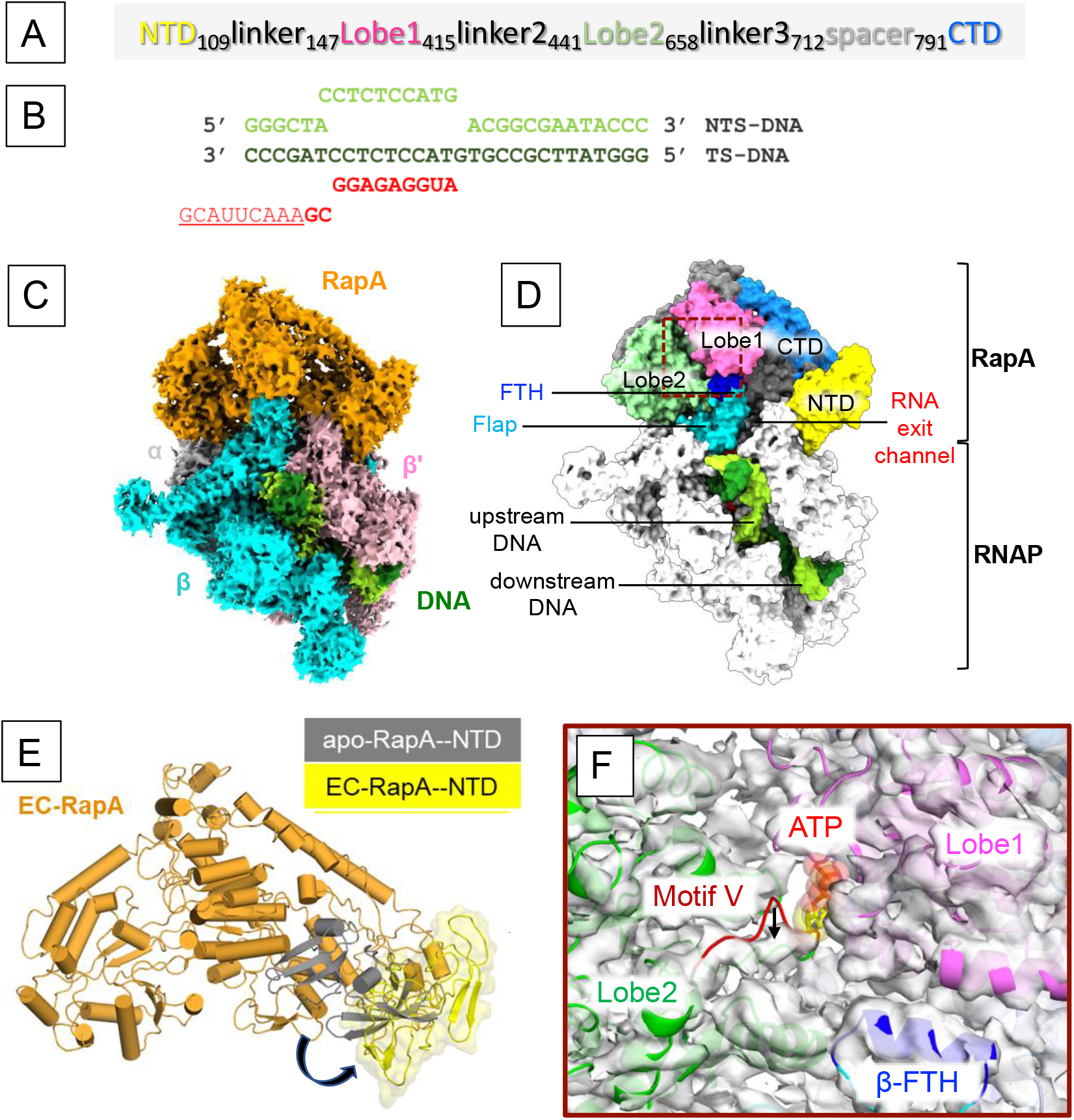
Cryo-EM structure of the RNAP-RapA elongation complex. **A)** The domain organization of RapA. **B)** The sequence of the nucleic acid scaffold used to form the elongation complex. **C)** Orthogonal view of the cryo-EM density map of EC-RapA. Subunits of RNAP and RapA are colored and labeled. Template and non-template DNA are shown in dark green and light green, respectively. **D)** Organization of EC-Rap. The structure of EC-RapA is depicted as a surface model. RapA and RNAP domains and are colored and labeled (FTH, flap-tip helix). The location of upstream and downstream DNA and the RNA exit channel are indicated. **E)** Movement of RapA-NTD upon binding to RNAP. RapA is depicted as a cartoon model. RapA-NTD in the EC-RapA (this study) and in the apo-form RapA (PDB: 6BOG) are colored yellow and dark gray, respectively. The shift of RapA-NTD is indicated by an arrow. **F)** Allosteric opening of the ATP binding site of RapA upon RNAP binding (brown dashed box in panel D). The apo-form RapA structure (cartoon model) fitted in the cryo-EM map of EC-RapA (white transparent). The Lobe domains of RapA are colored and labeled and the ATP binding is indicated by a modeled ATP. Compare with the apo-form RapA, motif V of the Lobe2 domain is shitted down in the EC-RapA complex as indicated by an arrow.

Biochemical studies of RapA characterized its enzymatic activities and modes of binding to RNAP (36-41). A model of RapA-mediated RNAP recycling proposes that RapA binds to a PTC, remodels it by utilizing its ATPase activity and releases the sequestered RNAP (35). The X-ray crystal structure of RapA revealed the organization of its ATPase catalytic domains: two RecA-like lobe domains and two Swi/Snf2-like domains together forming the ATP binding pocket (SFig. 1A) (42). The X-ray structure of the *E. coli* RNAP elongation complex (EC) with RapA (EC-RapA) revealed that RapA binds around the RNA exit channel of RNAP without conformational change from its apo-form structure (43) (SFig. 1D). This study speculated that RapA reactivates stalled RNAPs by means of an ATP-driven backward translocation mechanism. However, the role of RapA in RNAP recycling was not discussed.

To understand the mechanism of RNAP recycling by RapA, we determined four sets of cryo-electron microscopy (cryo-EM) structures including the *E. coli* RNAP EC, RNAP EC with RapA (EC-RapA), RNAP core enzyme (RNAP), and RNAP with RapA (RNAP-RapA binary complex). The structures reveal the conformational changes induced in RNAP and RapA upon their association. Based on the structural findings and the results of DNA binding assays, we propose that RapA functions in RNAP recycling by acting as a guardian; RapA prevents the non-specific association of the post-transcription terminated RNAP with DNA without obstructing σ factor binding to core enzyme (holoenzyme formation) and primes it for a next round of transcription cycle.

## RESULTS AND DISCUSSION

### Cryo-EM structure of the EC-RapA complex

EC-RapA was reconstituted *in vitro* by mixing RNAP with a DNA/RNA scaffold (Fig. 1B) to form the EC followed by adding RapA. The complex was cross-linked with BS3 and purified by gel filtration column. Cryo-EM data were collected after vitrifying purified complex supplemented with AMPPNP (a non-hydrolysable ATP analog) to capture ATP-bound state of RapA, and CMPCPP (a non-hydrolysable analog of CTP) to prevent RNAP backtracking. Unsupervised 3D classification of the particles in the processing steps revealed two distinct classes (SFig. 2) including EC-RapA (class2, 10%) and EC (class4, 36%). Bayesian polishing of particles gave final reconstructions of EC-RapA and EC at 3.3 Å and 3.15 Å resolutions, respectively.

**Figure 2.**
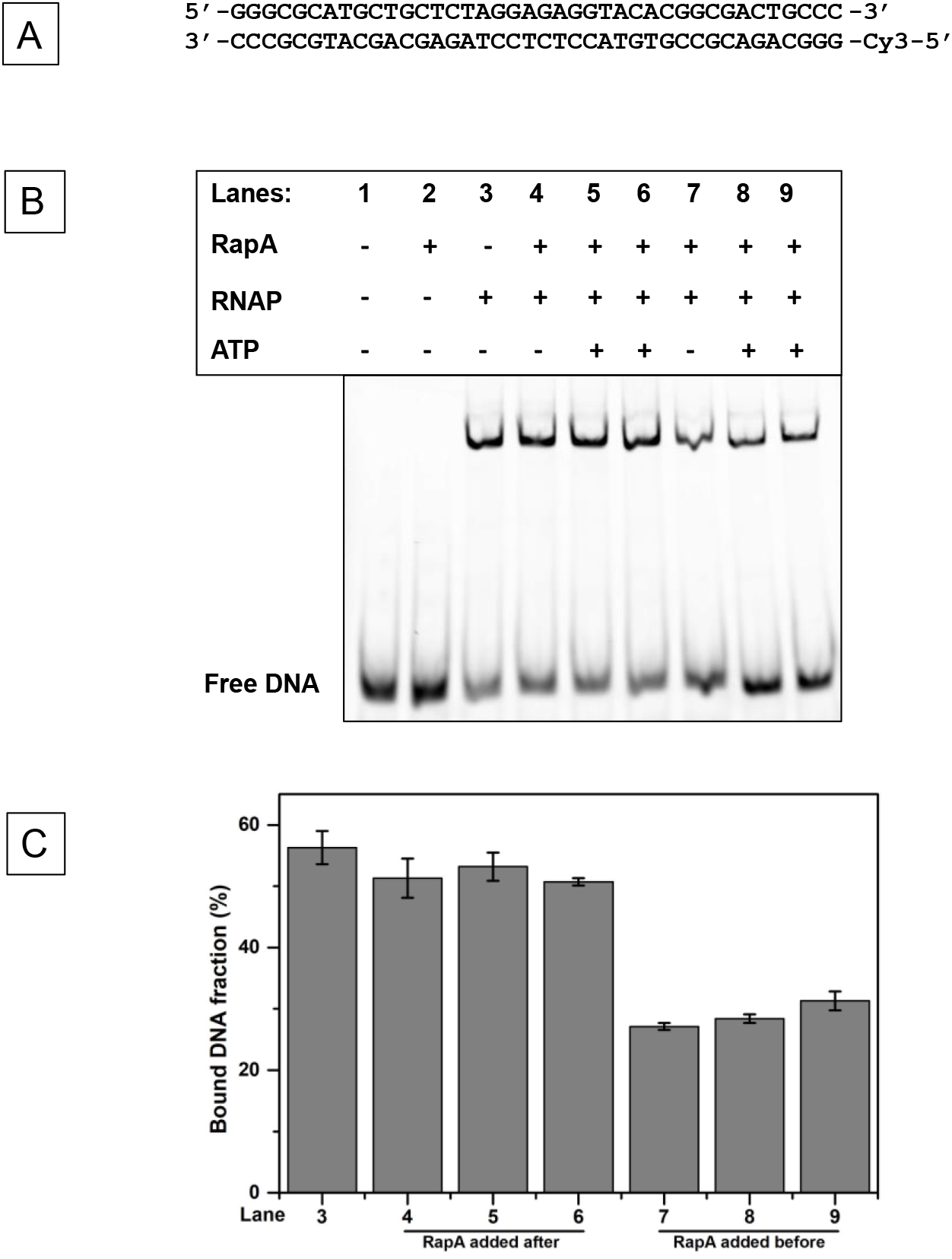
RapA reduces non-specific DNA binding of RNAP core enzyme. Electrophoretic Mobility Shift Assay (EMSA) to test the non-specific DNA binding to RNAP in the absence and presence of RapA. **A)** The sequence of Cy3-DNA used in the assay. **B)** RNAP was mixed with Cy3-labeled DNA and the shift DNA bands were quantitated. + indicated the components in the reaction mixture. In lanes 4-6, RapA was added to pre-formed RNAP-DNA complex. In lanes 7-9, DNA was added to pre-formed RNAP-RapA complex. In lanes 5 and 8, 1 mM ATP was added together with RapA. In lanes 6 and 9, RapA was p re-incubated with 1 mM ATP for 10 minutes before addition. **C)** Bar diagram showing the f raction of bound DNA (%) calculated with S.E., n=4.

The EC-RapA structure shows well-defined cryo-EM densities for RNAP (except ω subunit), RapA and the nucleic acid scaffold (Fig. 1C, SMovie 1). It also shows the density for CMPCPP at *i*+1 site of RNAP (SFig. 3A), but AMPPNP was not found in the RapA active site (SFig. 3B). RapA binds near the RNA exit channel of RNAP (Fig. 1D) as observed in the X-ray crystal structure of EC-RapA (SFigs. 1D) (43). Nonetheless, the cryo-EM structure of EC-RapA revealed the RNAP-induced conformational changes of RapA including the N-terminal domain (NTD, residues 1-109) and the ATP binding site compared to its apo-form (42) (Figs. 1E and F). The RapA-NTD rotates 90 degree and swings away from the Lobe 1 domain (Fig. 1E). The NTD rotation, together with the Lobe 1 domain creates a cavity that harbors the flap-tip helix (β subunit:897-905) and the zinc binding domain (ZBD, β’ subunit:70-88) of RNAP. The RapA-NTD rotation may influence the conformation of RapA active site allosterically; motif V of the Lobe 2 domain moves down toward a Lobe 2 helix (residues 540-553) compared with its position in the apo-form RapA (Fig. 1F). The spatial rearrangement of motif V, in turn, may facilitate better diffusion of ATP into the active site of RapA. NTD is involved in the regulation of ATPase activity of RapA. The ATPase activity of RapA derivative lacking NTD is five-fold higher than that of wild-type RapA. The ATPase activity of RapA is also enhanced upon binding to RNAP (44). The cryo-EM structure of EC-RapA elucidates a mechanism of enhancing ATPase activity of RapA upon binding of RNAP by which moving RapA-NTD from the Lobe 1 domain to widen the ATP binding site of RapA. Moreover, we propose a potential auto-inhibitory function of RapA-NTD; prior to binding RNAP, the RapA-NTD resides close to the catalytic lobe domains and may prevent non-specific ATP binding and hydrolysis.

**Figure 3.**
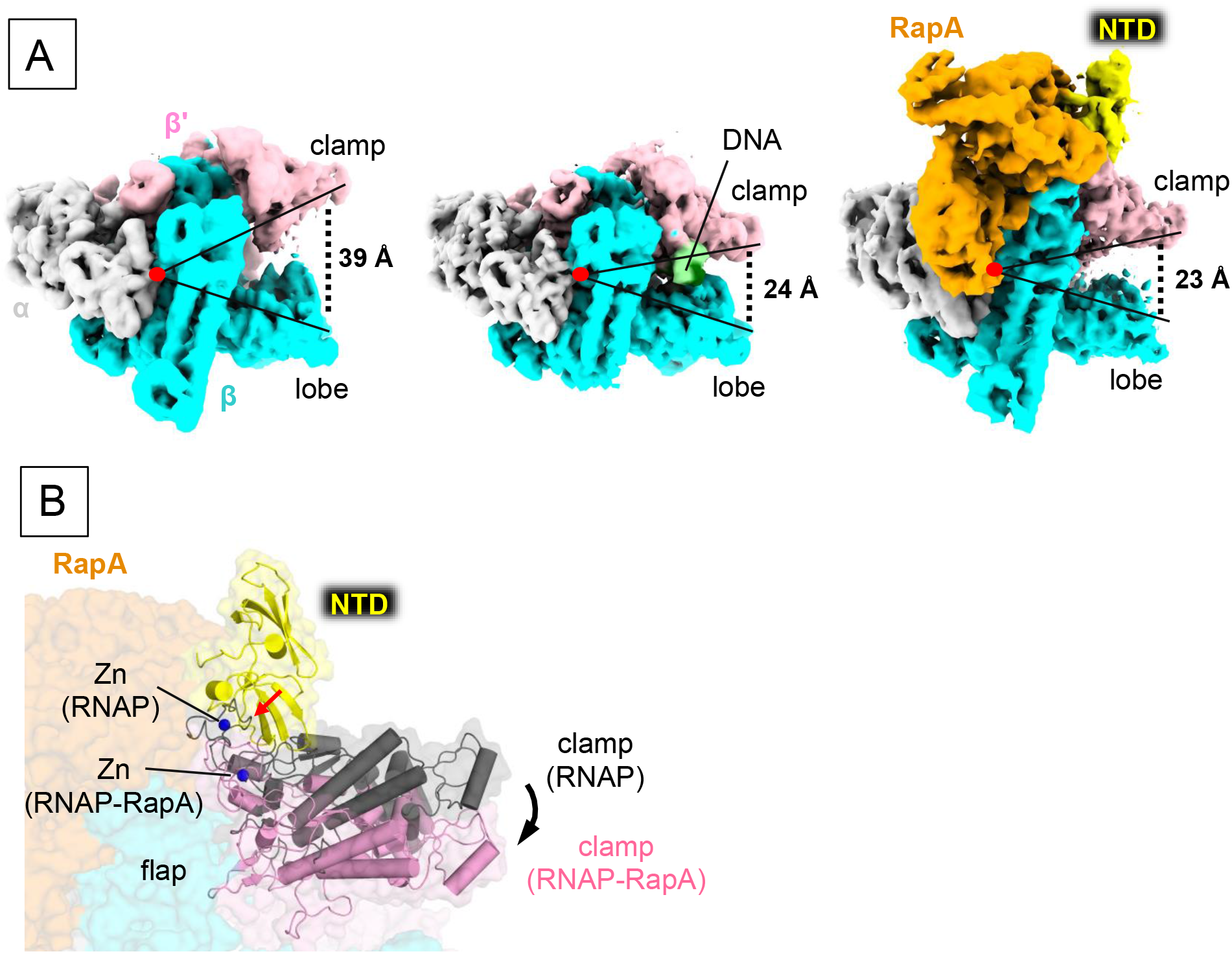
Cryo-EM structures of the RNAP core enzyme and the RNAP-RapA binary complex. **A)** Cryo-EM density maps of RNAP core enzyme (left), EC (middle) and RNAP-RapA (right). RNAP subunits and RapA are colored and labeled. RNAP domains and DNA are indicated. Opened and closed states of the RNAP clamp are represented by acute-angled lines and distances between the clamp and lobe domains of RNAP are indicated. **B)** RapA-mediated clamp closure. The RNAP-RapA is depicted as a transparent surface with cartoon models of the RapA-NTD and RNAP clamp as well as CPK representation of Zn bound at the ZBD. The clamp and Zn of apo-form RNAP are also depicted (gray). The structures are superimposed by aligning the DPBB domains of RNAP. Steric clash between the RapA-NTD and ZBD in the opened clamp is indicated by a red arrow and the RapA induced closing of the RNAP clamp is indicated by a black arrow.

### RapA constraints non-specific association of RNAP core enzyme with DNA

HelD in *Bacillus subtilis* and *Mycobacterium smegmatis* is the functional counterpart of RapA in *E. coli*; these ATP-dependent motor enzymes bind RNAP and facilitate RNAP recycling. Recent structural studies of the RNAP-HelD complex revealed that HelD associates with the main and the secondary channels of RNAP and opens the DNA binding main channel of EC to actively dissociate DNA/RNA from the EC (21,30,31). In sharp contrast, RapA accesses neither the main nor the secondary channels of RNAP but associates on the RNA exit channel of RNAP, and the RapA binding does not induce conformational change of EC, suggesting that the RapA does not dissociate DNA/RNA from the EC and it exploits an alternative mechanism to prevent the PTC formation and facilitates RNAP recycling.

A recent single-molecule study on transcription termination revealed that after RNA transcript release from EC, majority of RNAP remain bound on DNA and exhibit one-dimensional sliding over thousands of base pairs (45,46). Non-specific association of RNAP core enzyme with DNA hampers RNAP recycling and delays a next round of transcription. We, therefore, hypothesized that the RapA may function as a guardian of RNAP core enzyme for preventing the PTC formation by which reduces non-specific binding of RNAP with DNA.

To test this hypothesis, we performed electrophoretic mobility shift assay (EMSA) using a fluorophore (Cy3)-labeled DNA (Fig. 2A) and quantitated the fraction of DNA bound to RNAP. About 60 % of DNA was associated with RNAP in the absence of RapA (Figs. 2B and C). Addition of RapA to RNAP reduced the formation of RNAP-DNA complex by ∼2.5 fold. Supplementing ATP to RapA had no effect on the RNAP and DNA association. RapA did not reduce the RNAP-DNA complex if it was added to the pre-formed RNAP-DNA complex even in the presence of ATP. These results support our hypothesis that RapA reduces non-specific association of RNAP with DNA.

### Allosteric closure of RNAP clamp by RapA-NTD interaction with the zinc binding domain of RNAP

RNAP structure resembles a crab claw, with the β and β’ subunits forming two “pincers” that forms the main channel of RNAP serving the binding site for DNA (47). The pincer formed by the β’ subunit is also known as the “clamp”. Structural and biophysical studies of bacterial RNAP revealed flexible nature of the clamp, which can acquire different conformations from an “open” to a “closed” states (48,49). The clamp primarily remains in its “open” conformation in the apo-form RNAP core enzyme, but it closes upon formation of EC with DNA/RNA scaffold accommodated in the main channel. The clamp opening permits the entry of double-stranded DNA and/or DNA/RNA hybrid inside the main channel of RNAP. Although RapA binds around the RNA exit channel of RNAP, it may close the RNAP clamp allosterically and prevent non-specific DNA binding to RNAP. To study how the RapA influences the RNAP clamp conformation, we determined the cryo-EM structures of the RNAP core enzyme (SFig. 4) and RNAP-RapA binary complex (SFig. 5) at nominal resolutions of 3.41 Å and 4.8 Å, respectively.

**Figure 4.**
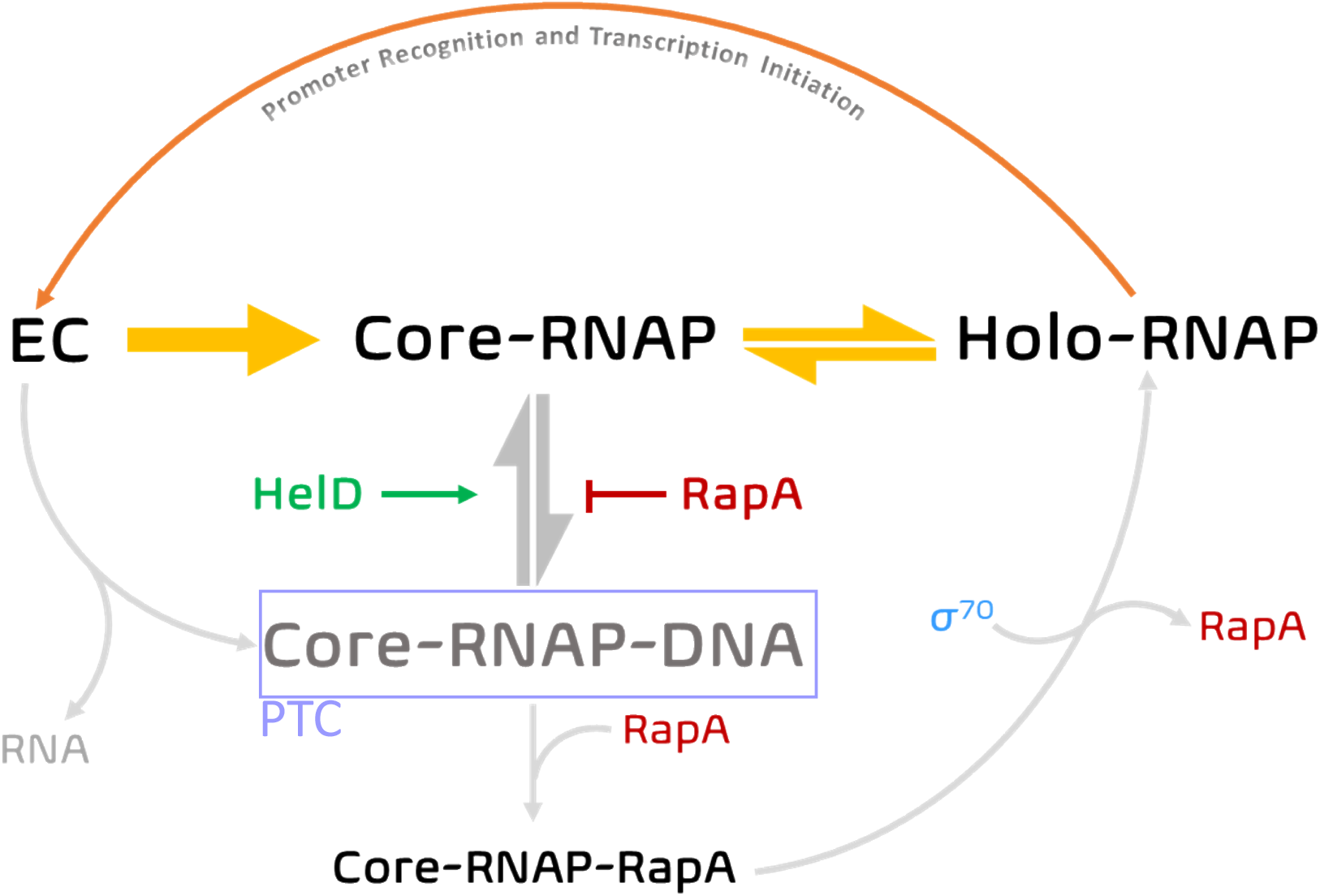
RapA-mediated RNAP recycling. Post-termination, RapA guards the free RNAP against non-specific association with DNA. RapA can also associate with RNAP released from a PTC. RapA is competed out from the RNAP-RapA binary complex upon σ factor recruitment and holoenzyme formation.

In the structure of RNAP core enzyme, the main channel is in an open conformation with the β’ clamp and β lobe being 39 Å apart (Fig. 3A, SMovie 2). In the structure of RNAP-RapA binary complex, the main channel is closed with the β’ clamp and β lobe distance of 23 Å (Fig. 3A, SMovie 2) as observed in the cryo-EM structures of EC (24 Å) and EC-RapA (24 Å) determined in this study.

The clamp closure of the RNAP-RapA binary complex is due to the positioning of RapA-NTD beside the zinc binding domain (ZBD) of the β’ subunit (Fig. 3B), which is a pivot point of the β’ clamp. Physical contact between RapA-NTD and β’-ZDB may restrict the flexibility of ZBD, and in turn, keeps the clamp in a closed state allosterically.

### Model for RapA-mediated RNAP recycling

The binding site of RapA on RNAP overlaps with that of the transcription elongation factor NusA (50) and are thought to be mutually exclusive (SFig. 6). Therefore, RapA is unable to access RNAP during the elongation phase of transcription. We posit that RapA prevents the detrimental effects of RNAP bound at non-specific DNA by acting as a recycling chaperone that binds RNAP core enzyme immediately after its release from DNA and RNA (Fig. 4). Upon RapA binding, it allosterically locks the RNAP in a closed clamp conformation and prevents its non-specific association with DNA. Moreover, as RapA binding site on RNAP does not mask the coiled-coiled (CC) domain of the β’ subunit of RNAP (Fig. 1D), the complex can still bind σ factor.

Both RapA and HelD enhance transcription by facilitating the RNAP recycling using distinct mechanisms. RapA recycles in a passive manner by allosterically closing the clamp of RNAP core enzyme, while HelD participates by interacting with the main channel of ECs, utilizing ATP energy and recycling RNAP. From bacterial genome sequence analysis, we found that presence of RapA and HelD is mutually exclusive with the former being predominantly present in proteobacteria whereas the latter is mostly present in fermicutes and actinobacteria (SFig. 7). This indicates that two different RNAP recycling mechanisms may have occurred during evolution. RapA cannot actively release DNA/RNA from EC, however, in *E. coli*, the transcription-repair coupling factor Mfd rescues EC stalled at DNA lesions and recruits the nucleotide excision repair (NER) machinery to the site of the damage (19). Binding of Mfd to the β subunit of stalled RNAP and upstream DNA of the transcription bubble (51,52) triggers its translocase activity (53). Mfd actively translocates along dsDNA and results in the collapse of the transcription bubble and dissociation of RNAP from the EC (54). Mfd-mediated active dissociation of RNAP may provide another safeguard to rescue RNAP that are bound non-specifically on DNA.

In summary, we propose that RapA functions as a guardian of RNAP after completion of RNA synthesis and releasing DNA from RNAP core enzyme, which is then protected by RapA for reducing its non-specific DNA binding. Binding of σ factor to the RNAP-RapA binary complex competes out RapA (42) and the new-formed holoenzyme is ready to begin the next round of transcription cycle.

## MATERIALS AND METHODS

### Protein purifications

*E. coli* RNAP core enzyme was overexpressed in *E. coli* BL21(DE3) cells transformed with pVS10 expression vector (encoding α, β, C-terminally His_6_-tagged β’ and ω subunits) (55) and grown in LB medium supplemented with ampicillin (100 µg/ml) at 37°C until OD_600_ = ∼0.5, and then added IPTG (final conc. 1 mM) and grown for 5 hours. RNAP enzyme was purified using Polymin P precipitation followed by heparin (HiTrap Heparin column), Ni-affinity (HisTrap HP column), and anion exchange (MonoQ column) chromatography steps (all columns from GE Healthcare). The purified RNAP core enzyme (20 µM) was suspended in the storage buffer (10 mM HEPES, pH 7.5, 50 mM NaCl, 0.1 mM EDTA, pH 8.0, 5 mM DTT), aliquoted, snap frozen in liquid N2, and stored at −80 °C.

*E. coli* RapA protein was overexpressed in *E. coli* BL21(DE3) cells transformed with pQE80L expression vector (encoding N-terminally His _6_-tagged full-length RapA) and grown in LB medium supplemented with ampicillin at 37°C till OD_600_=0.8. Then the RapA expression was induced with IPTG (final conc. 1 mM) at 37 °C for 3 hours and harvested (42). The protein was purified by affinity and size-exclusion chromatography using prepacked 5 ml Ni-affinity (HisTrap HP column), 5 ml heparin (HiTrap Heparin column), and Superdex200 columns (all columns from GE Healthcare). The purified RapA (230µM) in storage buffer (10 mM HEPES, pH 7.5, 50 mM NaCl, 0.1 mM EDTA, pH 8.0, 5 mM DTT) was aliquoted, flash frozen in liquid N2, and stored at −80 °C.

### Cryo-EM sample preparation

#### EC-RapA

The EC was reconstituted *in vitro* by mixing 4 µM RNAP core enzyme with equimolar amount of template DNA/RNA (Fig. 2A) in a storage buffer at 22 °C for 20 min, followed by mixing 6 µM non-template DNA for 10 min. The resulting EC was mixed with 5 µM RapA and incubated for 10 min at 22 °C. The EC-RapA complex was purified by a gel-filtration column (Superdex200, GE Healthcare) to remove excess DNA/RNA and RapA. The EC-RapA was concentrated to 4 mg/ml using Amicon Ultra centrifugal filter with 5kDa molecular weight cutoff (Merck Millipore). AMPPNP and CMPCPP (1 mM each) was added and incubated at 22 °C for 10 min, followed by BS3 (Sulfo DSS, 100µM) cross-linking for 30 minutes at RT and reaction was stopped by adding ammonium carbonate (1 M). The cross-linked EC-RapA was again passed through a gel-filtration column (Superdex 200, GE Healthcare) and concentrated to 2.5 mg/ml.

#### RNAP-RapA binary complex

The RNAP-RapA binary complex was assembled and purified in an identical fashion as the EC-RapA complex but omitting the nucleic acid scaffold and the nucleotide analogs. In brief, 4 µM RNAP was mixed with 5 µM RapA in a storage buffer and incubated for 10 min at 22 °C. The RNAP-RapA complex was pass through a gel-filtration column (Superdex200, GE Healthcare) to remove excess RapA. The RNAP-RapA complex was concentrated to 2.5 mg/ml using Amicon Ultra centrifugal filter with 5kDa molecular weight cutoff (Merck Millipore).

### Cryo-EM grids preparation

C-flat Cu grids (CF-1.2/1.3 400 mesh, Protochips, Morrisville, NC) were glow-discharged for 40 s prior to the application of 3.5 µl of the sample (2.5 –3.0 mg/ml protein concentration), and plunge-freezing in liquid ethane using a Vitrobot Mark IV (FEI, Hillsboro, OR) with 100 % chamber humidity at 5 °C.

### Cryo-EM data acquisition and processing

#### EC-RapA

The grids were imaged using a 300 kV Titan Krios (Thermo Fisher Scientific) equipped with a K2-Summit direct electron detector at the National Cryo-EM Facility (NCEF) at National Cancer Institute (NCI). A total of 1,674 movies were recorded with Latitude software (Gatan, Inc.) in counting mode with a pixel size of 1.32 Å and a defocus range of −1.0 to −2.5 µm. Data was collected with a dose of 4.64 e ^-^/s/physical pixel, with 15 sec exposure time (40 total frames) to give a total dose of 40 electrons/Å ^2^. Data was processed using RELION v3.0.8. Dose fractionated subframes were aligned and summed using MotionCor2 (56). The contrast transfer function was estimated for each summed image using Gctf (57). From the summed images, about 1,000 particles were manually picked and subjected to 2D classification in RELION (58). Projection averages of the most populated classes were used as templates for automated picking in RELION. Auto picked particles were manually inspected, then subjected to 2D classification in RELION specifying 100 classes. Poorly populated classes were removed, resulting in a total of 718,074 particles. 3D-classification of the particles was done in RELION using a model of *E. coli* core RNAP-RapA (PDB ID 4S20) low-pass filtered to 60 Å resolution using EMAN2 (59) as an initial 3D template. Among the 3D classes, the best-resolved classes (class2: 69,457 particles) and class4: 256,565 particles) were 3D auto-refined. Bayesian polishing and C TF refinement was performed on the particles and reconstructed maps were postprocessed in RELION (SFig. 2).

#### Core RNAP

The grids were imaged using a 300 kV Titan Krios (Thermo Fisher Scientific) equipped with a K3 Camera at NCI. A total of 3,072 movies were recorded with Latitude software (Gatan, Inc.) in counting mode with a pixel size of 1. 08 Å and a defocus range of −1.0 to −2.5 µm. Data was collected with a dose of 18 e^-^/s/physical pixel, with 3 sec exposure time (40 total frames) to give a total dose of 45 electrons/Å^2^. The date was processed using cryoSPARC (60). Dose fractionated subframes were aligned and summed using Patch-Motion correction job and the contrast transfer function was estimated for each summed image using Patch-CTF. A total of 766,796 particles were auto picked using Topaz picker (61), and then subjected to two rounds of 2D classification. Bad classes were removed, resulting in a total of 510,364 particles. Ab-initio model was generated, and the particles were subjected to multiple rounds of heterogeneous refinement. A final set of 235617 particles was refined and the reconstructed map was sharpened (SFig. 4).

#### RNAP-RapA binary complex

The grids were imaged using a 300 keV Titan Krios (Thermo Fisher Scientific) equipped with a K3 Camera (Gatan, Inc.) at NCI. A total of 3,504 movies were recorded with Latitude software (Gatan, Inc.) in counting mode with a pixel size of 1.08 Å and a defocus range of −1.0 to −2.5 µm. Data was collected with a total dose of 45 electrons/Å^2^. Movies were motion corrected using multi-patch motion correction and CTF values estimated using multi-patch CTF estimation in cryoSPARC (60). A total of 1,360,137 particles were auto picked using Topaz picker (61) and subjected to 2D classification job. Particles from selected good classes were used to generate initial models. All the particles were then further classified multiple times using heterogeneous refinement job-type to discard bad particles. A final set of 102,128 particles were then imported to RELION. To improve the density near the β lobe, focused classification was performed on RNAP density. The good class containing 88,511 particles was selected, and the particles were refined and post-processed (SFig. 5).

### Model building and refinement

To refine the EC-RapA complex structure, the crystal structure of EC-RapA (PDB: 4S20) was manually fit into the cryo-EM density map using Chimera (62), DNA and RNA were manually built by using Coot (63), and the initial model was real-space refined using Phenix (64). In the real-space refinement, the domains of RNAP and RapA were rigid-body refined and then subsequently refined with secondary structure, Ramachandran, rotamer and reference model restraints. The cryo-EM structures of the RNAP-RapA binary complex, RNAP core enzyme and RNAP-DNA-RapA complex were built using the cryo-EM structure of EC-RapA as a reference model, and these structures were refined as described in the EC-RapA complex structure refinement (STable 1).

### Electrophoretic Mobility Shift Assay

A linear Cy3-labeled dsDNA was generated by annealing the template and non-template strands by heating to 95 °C for 5 min and subsequent cooling to 10 °C at 1.5 °C/min (Fig. 3A). A final concentration of 150 nM Cy3-dsDNA was incubated with 300 nM RNAP for 10 minutes at 37 °C, followed by the addition of 300 nM RapA (either in the absence or presence of 1 mM ATP) and incubated for another 10 minutes (Fig. 3B, lanes 4-6). In the experimental set, first RNAP-RapA complex was formed by mixing 300 nM RNAP and 300 nM RapA (with/without 1 mM ATP), followed by incubation for 10 minutes at 37 °C. Then, 150 nM Cy3-dsDNA was added to the pre-formed complex, and further incubated for 10 minutes at 37 °C. Samples were loaded on a non-denaturing 4% polyacrylamide gel and electrophoresed in 0.5X TBE buffer. The Cy3-labeled DNA bands were visualized by a Typhoon imager and quantified using ImageQuant software (GE Healthcare).

## Data Availability

The cryo-EM density maps have been deposited in EMDataBank under accession codes EMDB: EMD-23900 (EC-RapA), EMD-23901 (EC), EMD-23902 (core RNAP) and EMD-23903 (RNAP-RapA). Atomic coordinates for the reported cryo-EM structures have been deposited with the Protein Data Bank under accession numbers 7KMN, 7KMO, 7KMP and 7KMQ.

## SUPPLEMENTARY DATA

This article contains supporting information.

## ACKNOWLEDGEMENTS

We thank Dr. Irina Artsimovitch at Ohio State University and Dr. Ding J. Jin at the National Cancer Institute/National Institutes of Health for providing the RNAP and RapA expression vectors, respectively. We thank Rishi K Vishwarkarma, Libor Krásný and Jan Dohnálek for valuable discussions and critical reading of this manuscript. We thank Jean-Paul Armache at Penn State for the technical support. We thank Carol Bator and Mike Carnegie at the Penn State Cryo-EM facility for supporting the cryo-EM data collections.

## FUNDING AND ADDITIONAL INFORMATION

This research was, in part, supported by the National Cancer Institute’s National Cryo-EM Facility at the Frederick National Laboratory for Cancer Research under contract HSSN261200800001E. This work was supported by NIH grants (R01 GM087350 and R35 GM131860 for K.S.M.).

---

**SFigure 1.**
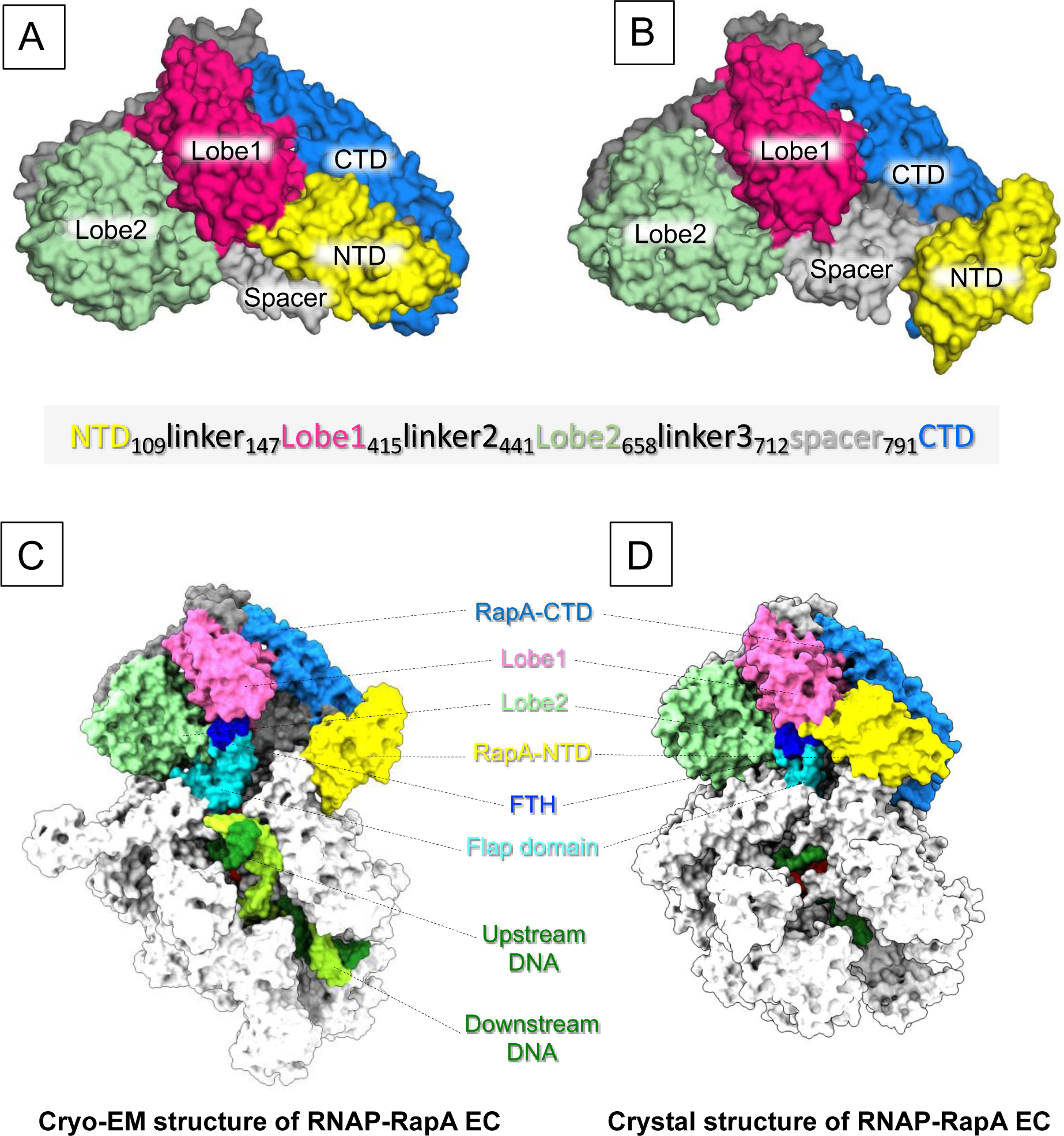
RapA conformation in apo-form and in RNAP bound form. RapA conformations found in the crystal structure of apo-form RapA (**A**) and in the cryo-EM structure of EC-RapA (**B**). Comparison of the RapA conformations in the cryo-EM structure (**C**) and the X-ray crystal structure of EC-RapA (**D**) (43).

**SFigure 2.**
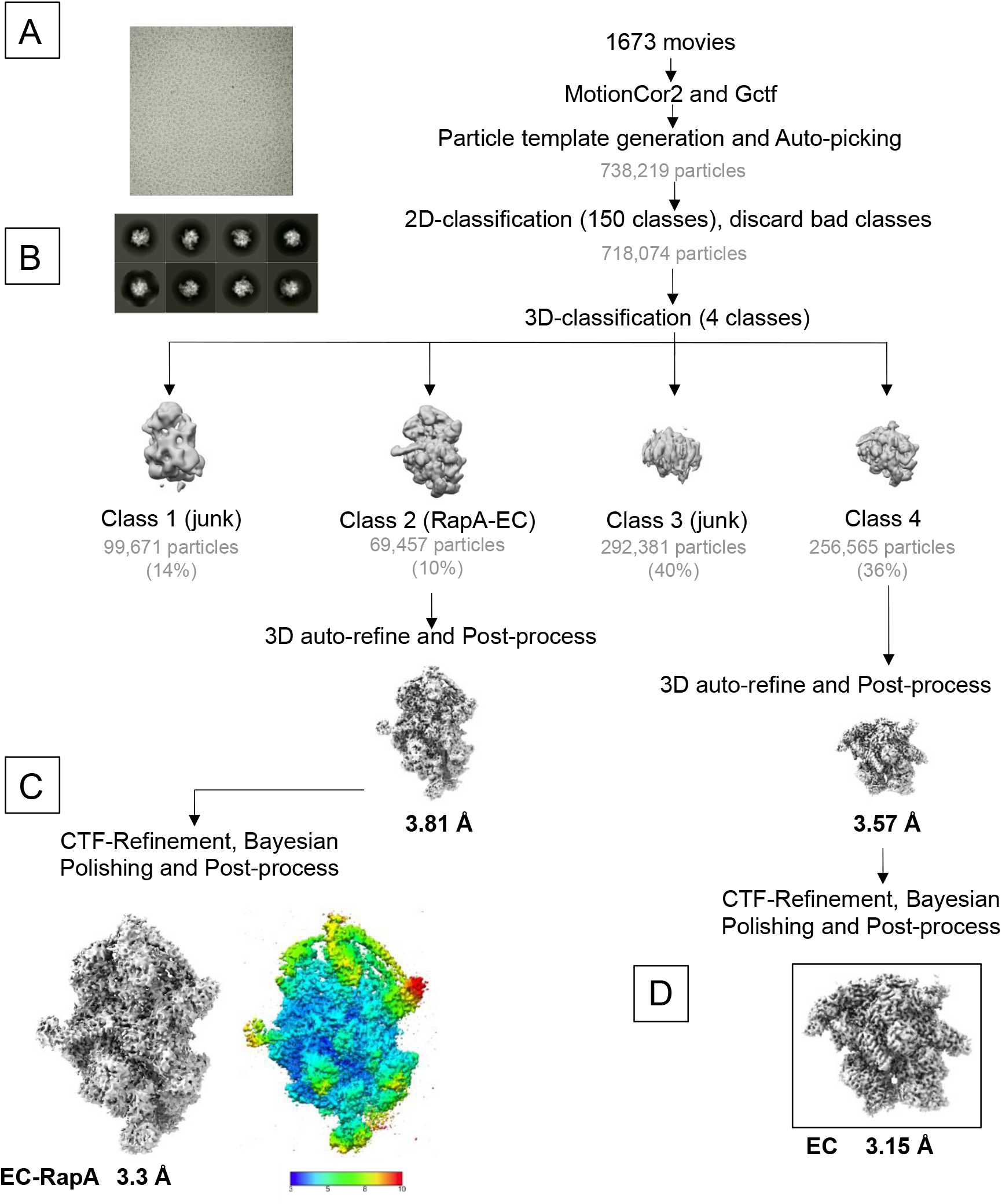
Cryo-EM processing pipeline for RNAP-RapA EC. **A)** A representative micrograph used for data processing. **B)** Selected representative 2D classes from 2D classification. **C)** The postprocessed cryo-EM density map of EC-RapA. The right view is identical to left but colored by local resolution. **D)** The postprocessed cryo-EM density map of the EC.

**SFigure 3.**
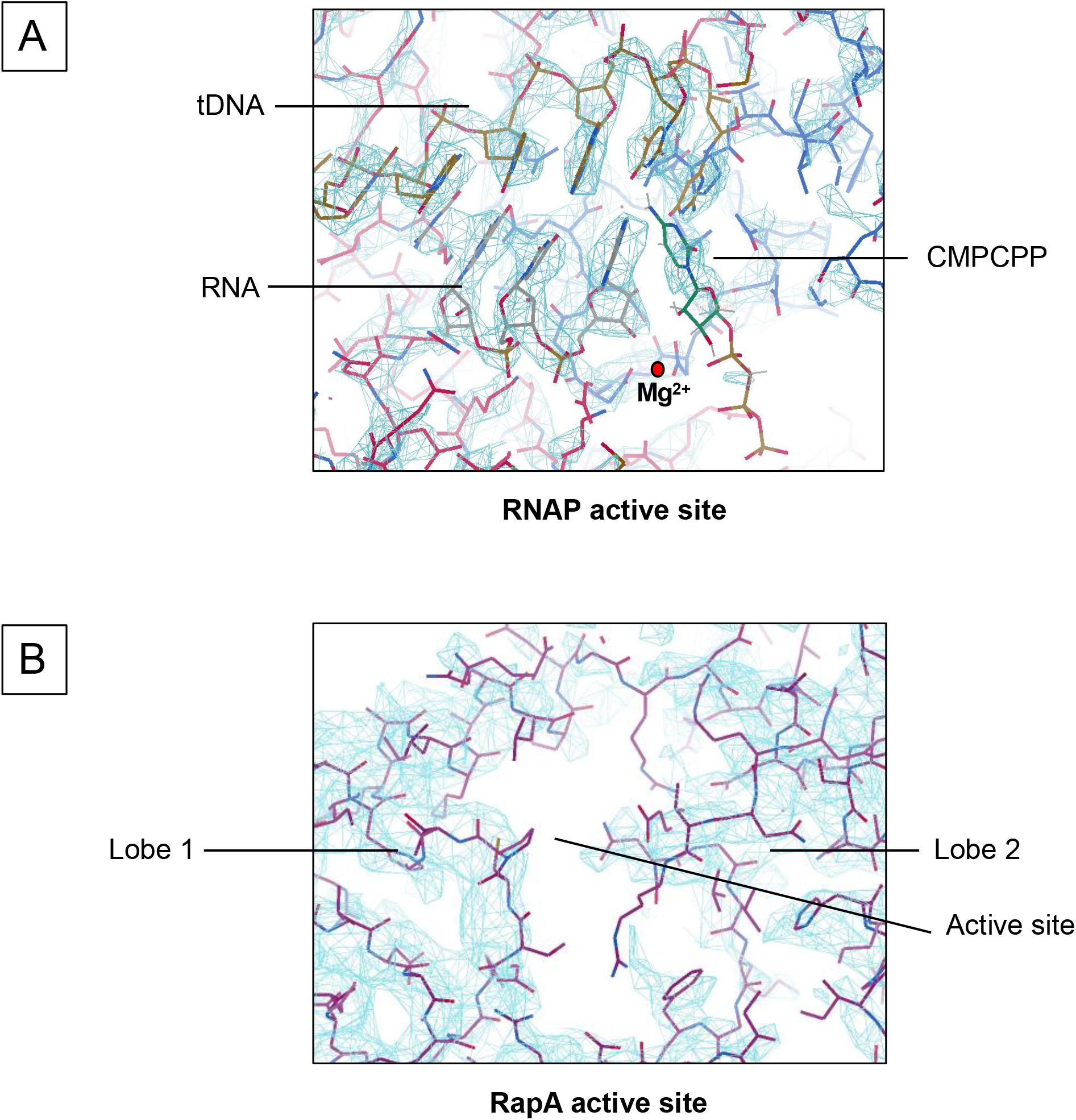
Cryo-EM density maps (blue mesh) at the active sites of RNAP (A) and RapA (B) in the EC-RapA.

**SFigure 4.**
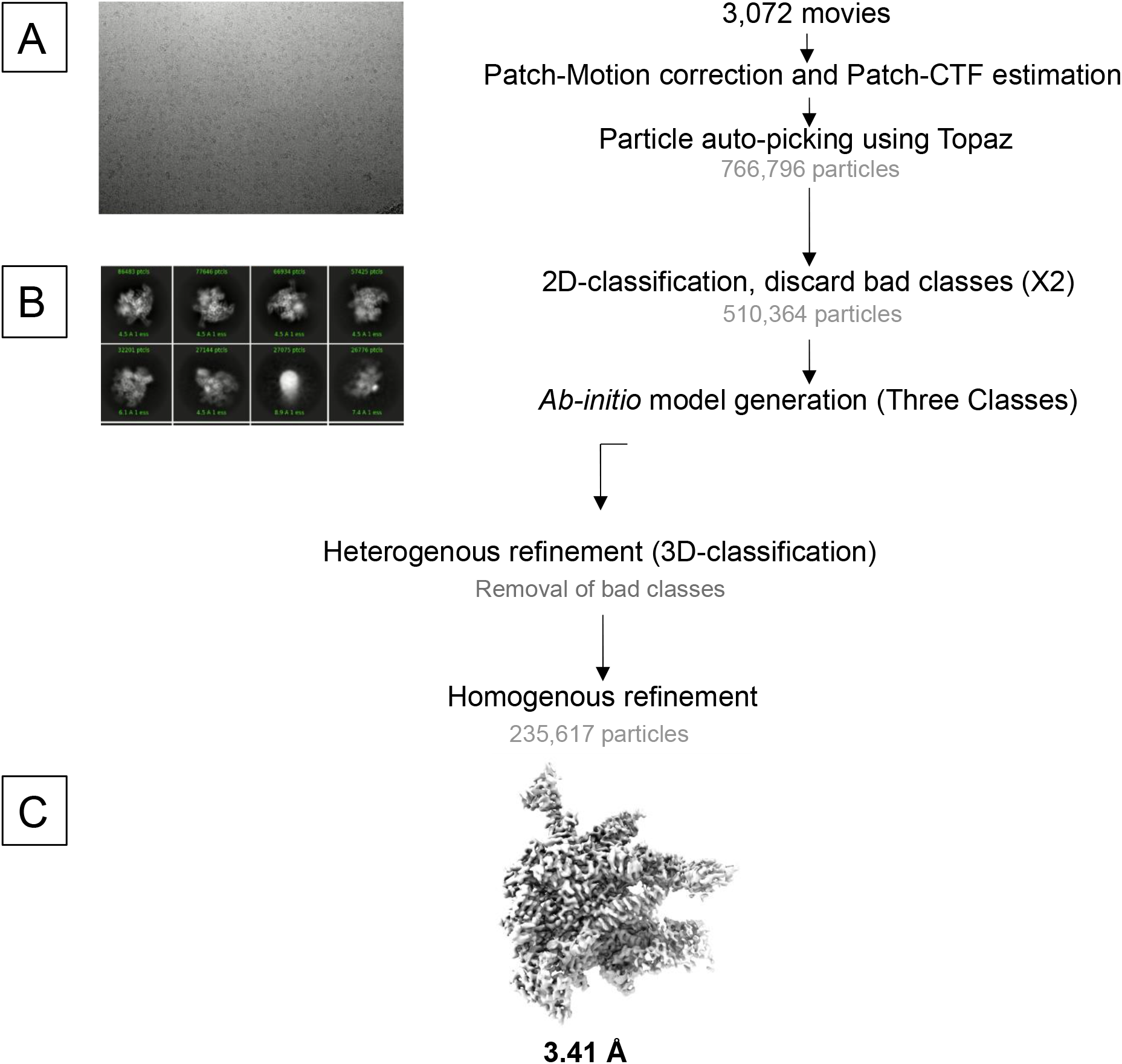
Cryo-EM processing pipeline for RNAP core enzyme. **A)** A representative micrograph used for data processing. **B)** Selected representative 2D classes from 2D classification. **C)** The postprocessed cryo-EM density map.

**SFigure 5.**
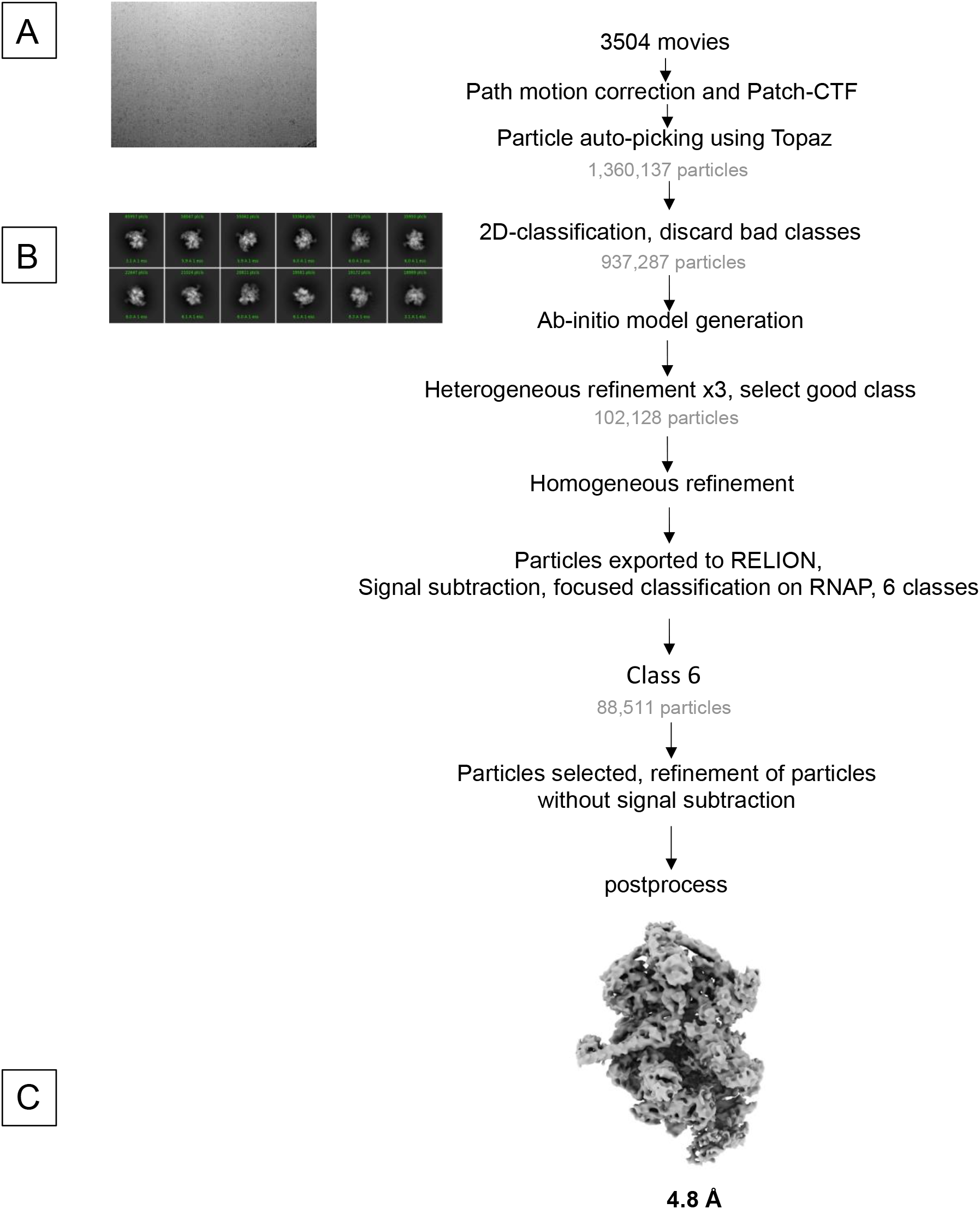
Cryo-EM processing pipeline for RNAP-RapA binary complex. **A)** A representative micrograph used for data processing. **B)** Selected representative 2D classes from 2D classification. **C)** The postprocessed cryo-EM density map.

**SFigure 6.**
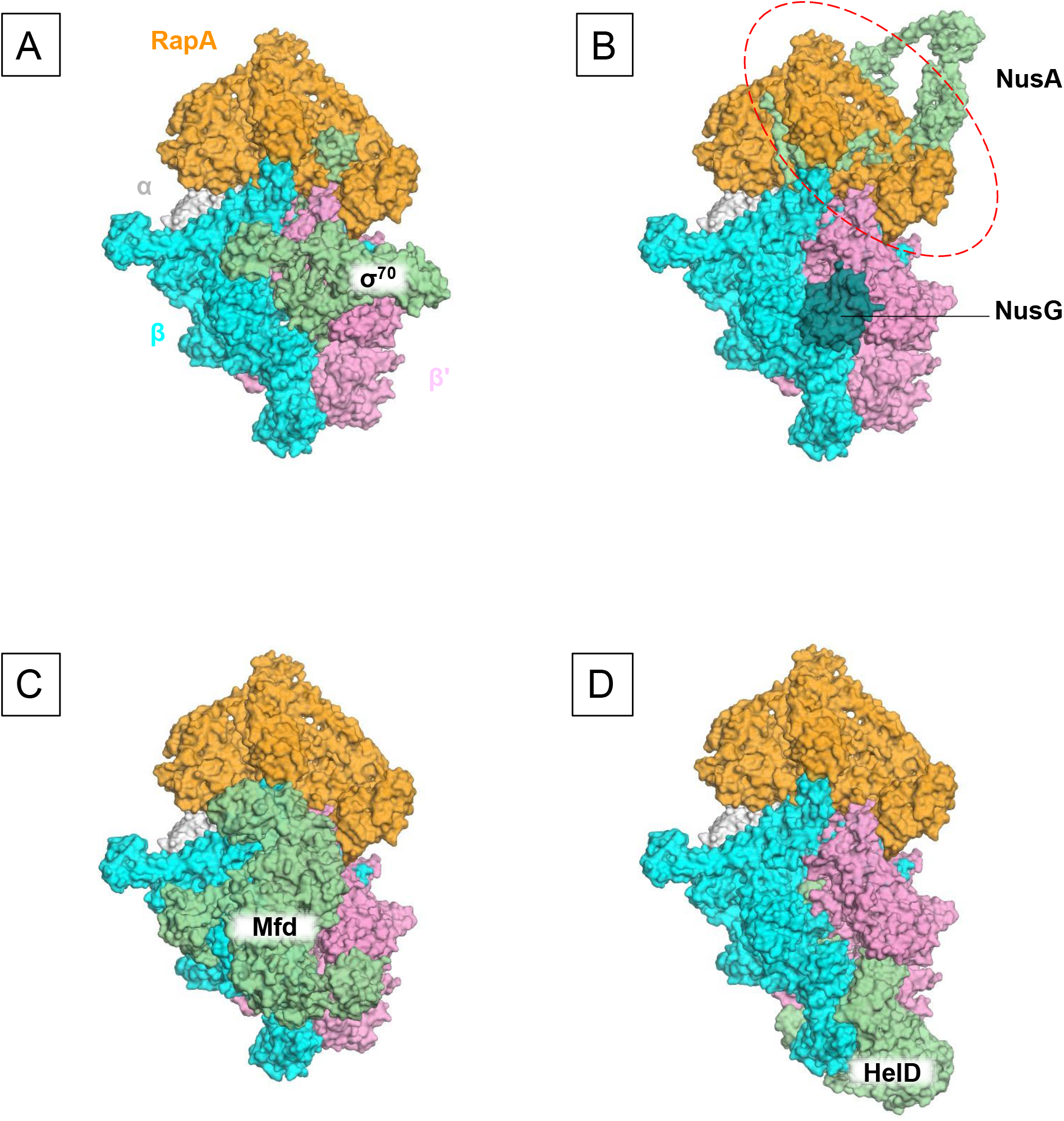
Binding sites of various transcription factors on RNAP compared to RapA. Occurrence of RapA and HelD is mutually exclusive. Panel (D) depicts them together to emphasize that their binding sites on RNAP are different.

**SMovie 1: Cryo-EM structure of *E. coli* RNAP elongation complex with RapA**

**SMovie 2: RapA-mediated allosteric closure of the RNAP clamp**

**Supplementary Table 1.** Cryo-EM data collection, refinement, and validation statistics.

|  | EC-RapA<br>(EMD-23900)<br>(7MKN) | EC<br>(EMD-23901)<br>(7MKO) | coreRNAP<br>(EMD-23902)<br>(7MKP) | RNAP-RapA<br>(EMD-23903)<br>(7MKQ) |
| --- | --- | --- | --- | --- |
| Data collection and processing |  |  |  |  |
| Magnification | 105,000 | 105,000 | 81,000 | 81,000 |
| Voltage (kV) | 300 | 300 | 300 | 300 |
| Electron exposure (e<sup>-</sup>/Å<sup>2</sup>) | 40 | 40 | 45 | 45 |
| Defocus range (μm) | -1.0 to -2.5 | -1.0 to -2.5 | -1.0 to -2.5 | -1.0 to -2.5 |
| Pixel size (Å) | 1.32 | 1.32 | 1.08 | 1.08 |
| Symmetry imposed | C1 | C1 | C1 | C1 |
| Initial particle images (no.) | 718,074 | 718,074 | 275,629 | 275,629 |
| Final particle images (no.) | 69,457 | 256,565 | 49,995 | 79,275 |
| Map resolution (Å) | 3.3 | 3.15 | 3.41 | 4.8 |
| FSC threshold | 0.143 | 0.143 | 0.143 | 0.143 |
| Map resolution range (Å) | 3.0-10.0 | 3.0-11.0 | 2.8-6.3 | 2.7-12.2 |
| Refinement |  |  |  |  |
| Initial model used (PDB code) | 4S20 | 7MKN | 7MKN | 7MKN |
| Model resolution (Å) | 4.7 | 3.3 | 3.3 | 3.3 |
| FSC threshold | 0.143 | 0.143 | 0.143 | 0.143 |
| *Model composition* |  |  |  |  |
| Non-hydrogen atoms | 34,084 | 26,269 | 24,943 | 32,822 |
| Protein residues | 4,177 | 3,194 | 2,986 | 4,182 |
| Ligands | Zn:2, Mg:1,<br>2TM | Zn:2, Mg:1<br>2TM | Zn:2, Mg:1 | Zn:2, Mg:1 |
| *B factors* (Å<sup>2</sup>) |  |  |  |  |
| Protein | 64.10 | 43.13 | 208.29 | 144.11 |
| Nucleotide | 109.24 | 84.86 | --- | --- |
| Ligand | 45.36 | 41.18 | 314.91 | 113.93 |
| *R.m.s. deviations* |  |  |  |  |
| Bond lengths (Å) | 0.005 | 0.007 | 0.010 | 0.004 |
| Bond angles (°) | 1.004 | 1.048 | 1.212 | 0.910 |
| *Validation* |  |  |  |  |
| MolProbity score | 2.3 | 2.33 | 2.34 | 2.18 |
| Clash score | 21.57 | 20.99 | 21.88 | 16.50 |
| Rotamer outliers (%) | 0.90 | 1.22 | 0.52 | 1.16 |
| *Ramachandran plot* |  |  |  |  |
| Favored (%) | 92.50 | 91.60 | 91.52 | 92.58 |
| Allowed (%) | 7.02 | 7.93 | 8.07 | 6.99 |
| Disallowed (%) | 0.48 | 0.47 | 0.41 | 0.43 |
| Model vs. Data |  |  |  |  |
| CC (mask) | 0.69 | 0.75 | 0.80 | 0.79 |
| CC (box) | 0.71 | 0.71 | 0.90 | 0.81 |
| CC (peak) | 0.59 | 0.61 | 0.77 | 0.71 |
| CC (volume) | 0.69 | 0.72 | 0.80 | 0.79 |
| Mean CC for ligands | 0.68 | 0.71 | 0.81 | 0.85 |

